# Prenatal Maternal Inflammation Is Associated with Altered Offspring Mesolimbic White Matter Circuitry Observed in Late Midlife

**DOI:** 10.64898/2026.04.06.716489

**Authors:** Ranesh Mopuru, Blake Elliott, Linda J. Hoffman, Naoya Tani, Ann M. Kring, Elizabeth C. Breen, Barbara A. Cohn, Piera M. Cirillo, Nickilou Y. Krigbaum, Mark D’Esposito, Ashby B. Cogan, Bhakti P. Patwardhan, Thomas Olino, Ingrid R. Olson, Lauren M. Ellman

## Abstract

**Background:** Exposure to prenatal maternal inflammation (PNMI) has been linked to neurodevelopmental alterations in human offspring. Preclinical studies suggest that PNMI disrupts reward circuitry, particularly within mesolimbic circuits. However, the effects of PNMI on mesolimbic circuits (i.e, ventral tegmental area (VTA) projections to the hippocampus (VTA-H) and limbic striatum (VTA-LS)) in humans are not yet known.

**Methods:** Data for PNMI biomarkers [interleukin (IL)-6, IL-8, IL-1 receptor antagonist (IL-1ra), soluble TNF receptor-II (sTNF-RII)] from first trimester (T1) and second trimester (T2) maternal sera, and offspring MRI brain scans in late midlife (aged 57–63 years), were available for 89 mother–offspring dyads. Probabilistic tractography delineated bilateral VTA-H and VTA-LS tracts. Macrostructural tract measures were examined using hierarchical linear regressions. Microstructural integrity was assessed using neurite orientation dispersion and density imaging, and permutation-based cluster analyses.

**Results:** Higher T2 IL-1ra was associated with increased macrostructure (left VTA–H tract), whereas higher T2 sTNF-RII was associated with reduced macrostructure (right VTA–H and VTA–LS tracts) and higher T2 IL-8 (bilateral VTA-LS tracts). Microstructurally, higher T2 IL-6 was associated with increased neurite density (distal cluster, right VTA–H tract), while higher T1 IL-8 was associated with reduced neurite density (near the hippocampus in the left VTA–H tract, near the VTA in bilateral VTA–LS tracts).

**Conclusions:** PNMI was associated with altered mesolimbic reward circuitry in offspring. This suggests that prenatal inflammation may contribute to affective and motivational disorders in offspring via alterations in mesolimbic circuitry.

## Introduction

The prenatal period is a critical phase of neurodevelopment, where perturbations to the intrauterine environment can be associated with impaired neurological structure and function that may influence the health trajectories of offspring (1–3). Inflammation is a key regulator of the in utero environment and is tightly regulated in a healthy pregnancy to ensure optimal outcomes for both the offspring and the mother (4). Deviations from the homeostatic immunological state in pregnancy is associated with various neuropsychiatric disorders in offspring such as autism spectrum disorder, attention-deficit hyperactivity disorder, depression, and schizophrenia (5–8).

One possible biological mechanism linking inflammation to these disorders is alterations in offspring brain structure and function caused by prenatal maternal inflammation (PNMI) (9–14). Pertinently, the effects of PNMI persist into adulthood, with structural and functional alterations being observed in various regions, including limbic regions (10,15,16). For example, elevated PNMI is associated with lower volumes in the entorhinal cortex and posterior cingulate in schizophrenia spectrum offspring, whereas no significant associations between PNMI and structural brain changes were observed in healthy control offspring (10). In a separate cohort oversampled for obstetric complications, elevated PNMI was associated with disrupted hippocampal and hypothalamic connectivity and activity (15). Recent work from our lab has shown that elevated PNMI was associated with changes in hippocampal gray matter in a community-based cohort of late-midlife offspring. Specifically, there was lower neurite density within hippocampal subfields CA3, CA4, dentate gyrus, and subiculum (16). Importantly, many of the limbic regions shown to be sensitive to PNMI are also integral components of the brain’s dopaminergic circuitry. In particular, portions of the hippocampus regulate dopamine neuron activity in the ventral tegmental area (VTA) (17,18). However, the direct effect of PNMI on classic reward circuits in humans has yet to be studied.

Dopamine is a critical neuromodulator implicated in motivation, learning, memory, and decision making (19–22). Impaired dopaminergic signaling is implicated in the neuropsychiatric disorders associated with higher PNMI (23–26), contributing to dysfunction in symptom domains such as anhedonia, avolition, impulsivity, and novelty-seeking (27–30). Evidence from preclinical studies has shown that PNMI disrupts the development of dopaminergic pathways in offspring, leading to long-lasting alterations in dopamine receptor expression, dopamine levels, and tyrosine hydroxylase expression (31,32). These disruptions may underlie the increased risk for neuropsychiatric disorders associated with PNMI, particularly those with symptoms linked to dopamine dysregulation (e.g., motivational and reward-processing deficits). Despite strong evidence from animal models implicating PNMI-related alterations in dopaminergic circuitry, the structural organization of mesolimbic dopamine pathways in humans, and their sensitivity to prenatal inflammatory exposure, remains poorly understood.

The ventral tegmental area (VTA) is a midbrain nucleus that produces dopamine and hence, is a critical hub of the dopamine system. The VTA has primarily been studied for its central role in motivation and reward processing through its efferent dopaminergic projections to limbic and cortical structures through the mesolimbic and mesocortical pathways (33,34). The mesolimbic pathway consists of VTA projections to the limbic striatum, implicated in reward and motivated behavior (35,36), as well as direct dopaminergic projections to the hippocampus, as part of a larger VTA-Hippocampus loop, which plays a role in motivated memory encoding (18,37,38). Although preclinical models have provided insights into mesolimbic VTA circuitry, research in humans, particularly on the effects of PNMI on these pathways, remains limited.

Given evidence that PNMI is linked to alterations in limbic brain regions that regulate dopamine signaling preclinically, and that dopaminergic dysfunction is a hallmark of several PNMI-associated psychiatric outcomes, characterizing the integrity of mesolimbic pathways in humans is a critical next step. The current study investigated the effects of PNMI on human mesolimbic structural connectivity in vivo using probabilistic tractography (macrostructure) and microstructural modelling techniques. Combining these complementary tract metrics (macrostructure and microstructure) allow us to disentangle how PNMI affects both the large-scale architecture of white matter pathways and their internal, voxel-wise neurite microstructure. Macrostructure may reflect the overall tract geometry and communication capacity of each tract (39–41) and microstructural indices provide an index of myelination and microscopic changes in the processes that extend from neurons (40,42,43) with changes in neurite density reflecting disrupted axonal integrity within the mesolimbic white matter pathways.

Based on prior preclinical and human evidence linking PNMI to limbic structural alterations and dopaminergic dysfunction, we tested four related hypotheses. First, we hypothesized that higher PNMI would be associated with reduced tract macrostructure in mesolimbic pathways connecting the VTA with the hippocampus and limbic striatum, reflecting altered large-scale pathway organization. Second, we hypothesized that elevated PNMI would be associated with reduced tract microstructure, indicative of compromised neurite integrity. Third, based on previous findings of structural changes in limbic regions due to PNMI exposure, we predicted the effects would be specific to interleukin-6 (IL-6), IL-8, and IL-1 receptor antagonist (IL-1ra) (10,16). Finally, given prior findings of PNMI-related microstructural alterations within hippocampal gray matter (16), we predicted that associations between PNMI and microstructure would be strongest in the distal, hippocampal portions of the VTA-hippocampus tract.

## Methods

### Participants

Participants were drawn from the Healthy Brains Project (HBP), a late midlife follow-up study of the Child Health and Development Studies (CHDS), a prospective birth cohort of 19,044 live births to pregnant women receiving care through the Kaiser Foundation Health Plan in Alameda County, California between 1959 and 1966. A subset of offspring participated in the CHDS adolescent study (n=2,020) and archived first (T1) and/or second (T2) trimester maternal serum samples were later assayed for PNMI biomarkers in 737 dyads.

The HBP recruited a subset of these offspring in late midlife using a tiered strategy prioritizing individuals with existing PNMI data and feasible proximity to the study site. The current analyses drew from 139 HBP participants, of whom 101 had diffusion MRI data and T1 or T2 biomarker data. Following visual quality inspections of MRI data, 89 participants had usable scans. The final sample included n=82 with usable MRI and T1 sera data and n=80 with MRI and T2 data (see Supplementary Figure 1). All procedures were approved by Institutional Review Boards (IRBs) at Temple University, University of California Berkeley, and the Public Health Institute of California. All HBP participants provided written informed consent, and because CHDS predated IRBs, mothers voluntarily provided verbal permission for access to medical records for themselves and their children.

### Demographic Data

Sociodemographic data was collected for mothers primarily during the second trimester of pregnancy (mean gestation = 15.3 weeks, SD = 6.8 weeks). Maternal education was chosen as a proxy for socioeconomic status (SES) since it had the most complete data of all available SES measures collected and has been shown to be correlated with other measures of SES such as annual household income, in similar published studies (7,44,45). Maternal education was classified into three categories “Did not complete High School”, “Completed High School”, or “Completed more than High School” and coded as a factor variable for analyses. Offspring sex assigned at birth (referred to as “sex” throughout the paper), was categorized as “Male” or “Female”. Detailed demographic information is provided in Table 1.

**Table 1.** Participant Demographics.

|  |  |
| --- | --- |
| **Offspring Age** (years) Mean (SD) [range] | 59.6 (1.29) [57.3-62.3] |
| **Offspring Sex** % (n) Male | 46.1 (41) |
| **Offspring Ethnicity** % (n) Hispanic | 7.2 (6) |
| **Offspring Race** % (n) |  |
| American Indian/Alaska Native | 5.6 (5) |
| Asian | 11.2 (10) |
| Black | 14.6 (13) |
| White | 75.3 (67) |
| More than one race | 9.0 (8) |
| Unknown | 1.1 (1) |
| **Maternal Education** % (n) |  |
| < High school | 7.9 (7) |
| = High school | 37.1 (33) |
| > High school | 55.1 (49) |

Note. N = 89; demographics are from all mother-offspring dyads who were included in at least one analysis.

### Prenatal Maternal Inflammation (PNMI) Data

Data were available for PNMI biomarker concentrations determined in archived maternal sera from a subset of pregnant mothers in the parent CHDS study (n=737), during T1 (mean gestational age in weeks 12.1 ± 2.6) and/or T2 (mean gestational age in weeks 23.7 ± 3.0), as previously described (7,44,45). The biomarkers were interleukin-6 (IL-6), IL-8, IL-1 receptor antagonist (IL-1ra), and soluble tumor necrosis factor receptor-II (sTNF-RII).

### NODDI Metrics

Detailed MRI methods are available in the supplementary information. The NODDI model was fitted to participants’ preprocessed diffusion images using the AMICO pipeline Version 1.4.2 to produce estimates of neurite density for the whole brain (46). The NODDI model characterizes three separate types of water diffusion into unique microstructural compartments: intracellular volume fraction is modelled as neurite density index (NDI), the orientation or dispersion of neurites modelled as orientation dispersion index (ODI), and isotropic diffusion modelled as the free water fraction (FWF) (43). We only examined NDI both to limit the number of models in our analyses, and because it is generally regarded as the most sensitive NODDI metric for white matter microstructure (47).

### Probabilistic Tractography

Probabilistic tractography was performed in MRTrix3 (https://www.mrtrix.org/) using the iFOD2 algorithm with the *tckgen* command in subjects’ native space. All tractographies seeded in the VTA and were performed unidirectionally, with the upstream target (hippocampus or limbic striatum) designated as both the inclusion and stopping criteria. To allow inferences about tract density (macrostructure), we seeded a fixed number of random streamline attempts per voxel in the seed mask. To control for individual variation in seed size, the lowest volumetric occurrence of the VTA within the cohort was determined. We found the smallest size of the VTA in any subject to have a volume of 276 voxels. Using 5,000 seeding attempts per voxel as the default selection criteria, we calculated the total number of viable seeds for all tractographies by multiplying 276 voxels by 5,000 seed attempts per voxels, resulting in 1,380,000 total seeds for each tractography; an approach that was similar to prior tractography work (48). For each tract, we excluded either the left or right hemisphere to ensure we only captured streamlines that projected to the target region ipsilaterally for all subjects.

### Statistical analyses

All analyses were performed in R (Version 4.3.0). Due to substantial skew, all PNMI biomarker values were log-transformed prior to analysis, consistent with standard practice for these markers (7,44,45). Furthermore, tract macrostructure values were also log-transformed to address non-normal distribution.

Spearman’s correlations were used to assess bivariate relationships between PNMI biomarkers and tract macrostructure. Maternal education and intracranial volume (ICV) were included as covariates in models to control for socioeconomic status and structural variation in brain size. Of the other potential covariates examined (offspring sex, age, and handedness), none were significantly associated with one independent or dependent variable (see supplementary Tables 1 and 2). As such, only maternal education and ICV were retained as covariates in the subsequent analyses.

**Table 2.** Results of hierarchical linear regression analyses for prenatal maternal inflammation and offspring tract streamline density (Bootstrapped for 5,000 iterations)

|  |  | Left VTA-Hippocampus |  | Right VTA-Hippocampus |  | Left VTA-Limbic Striatum |  | Right VTA-Limbic Striatum |  |
| --- | --- | --- | --- | --- | --- | --- | --- | --- | --- |
| Independent Variable | | *b* (C.I) | $\Delta R^2$ | *b* (C.I) | $\Delta R^2$ | *b* (C.I) | $\Delta R^2$ | *b* (C.I) | $\Delta R^2$ |
| **Step 1: Covariates** |  |  | 0.002 |  | 0.061 |  | 0.111 |  | 0.080 |
| Intracranial Volume |  | 1.99e-7<br>[-8.3e-7,1.2e-6] |  | 3.60e-7<br>[-5.8e-7,1.3e-6] |  | -1.34e-6<br>[-2.9e-6,2.6e-7] |  | -1.07e-6<br>[-2.6e-6,5.1e-7] |  |
| Maternal Education = High School |  | 0.045<br>[-0.72,0.81] |  | -0.388<br>[-1.08,0.31] |  | -0.149<br>[-1.03,1.33] |  | -0.036<br>[-1.21,1.13] |  |
| Maternal Education > High School |  | -0.005<br>[-0.75,0.74] |  | **-0.677**<br>**[-1.35,-3.4e-3]\*** |  | 1.009<br>[-0.14,2.16] |  | 0.704<br>[-0.43,1.84] |  |
| **Step 2: Additive Effect** |  |  |  |  |  |  |  |  |  |
| **First Trimester** | IL-1ra | -0.066<br>[-0.43,0.37] | 0.016 | 0.090<br>[-0.22,0.40] | 0.010 | -0.018<br>[-0.72,0.70] | 0.016 | -0.598<br>[-1.42,0.22] | 0.072 |
|  | IL-6 | -0.004<br>[-0.48,0.29] | 0.010 | -0.275<br>[-0.65,0.02] | 0.033 | -0.196<br>[-0.90,0.42] | 0.015 | -0.429<br>[-1.10,0.11] | 0.034 |
|  | IL-8 | 0.021<br>[-0.12,0.15] | 0.013 | -0.075<br>[-0.19,0.04] | 0.030 | -0.117<br>[-0.30,0.08] | 0.023 | -0.127<br>[-0.30,0.05] | 0.028 |
|  | <u>sTNF-RII</u> | -0.179<br>[-0.80,0.63] | 0.011 | **-0.671**<br>**[-1.37,-0.02]\*** | **0.049** | -0.066<br>[-1.16,0.97] | 0.007 | **-1.226**<br>**[-2.71,-0.24]\*** | **0.070** |
| **Second Trimester** | IL-1ra | **0.416**<br>**[0.02,0.90]\*** | **0.046** | 0.185<br>[-0.16,0.57] | 0.017 | -0.108<br>[-1.06,0.84] | 0.020 | -0.329<br>[-1.13,0.43] | 0.022 |
|  | IL-6 | 0.174<br>[-0.05,0.39] | 0.029 | -0.028<br>[-0.27,0.15] | 0.007 | -0.390<br>[-0.74, 0.15] | 0.042 | -0.205<br>[-0.48,0.06] | 0.017 |
|  | IL-8 | 0.106<br>[-0.03,0.23] | 0.035 | -0.007<br>[-0.18,0.16] | 0.014 | **-0.248**<br>**[-0.49,-0.01]\*** | **0.055** | **-0.247**<br>**[-0.47,-0.06]\*** | **0.061** |
|  | <u>sTNF-RII</u> | -0.192<br>[-1.13,0.69] | 0.014 | -0.316<br>[-1.03,0.37] | 0.017 | -0.470<br>[-1.83,0.72] | 0.015 | -0.156<br>[-1.67,1.10] | 0.013 |

Note. Each of the additive effects were assessed using separate models; IL = Interleukin, RA = receptor antagonist, sTNF-RII = soluble tumor necrosis factor receptor-II, \* $p < 0.05$ .

### Tract Macrostructure

Streamline counts of each tractography, controlling for seed size, were used as an index of tract macrostructure. Following previous research (7,16) separate hierarchical linear regressions were performed to examine the association between T1 and T2 inflammatory biomarkers and tract macrostructure in both hemispheres of the VTA-hippocampus and VTA-limbic striatum tracts. In each model, ICV and maternal education were entered in the first step to account for structural and socioeconomic variance. PNMI biomarkers were added in the second step in separate models to assess their contribution to tract macrostructure. Given the moderate sample size and our anticipated small-moderate effect sizes based on prior findings in the PNMI literature (10,11,15,16), all models were bootstrapped using 5,000 iterations to improve estimate stability. Bootstrapped p-values and confidence intervals were interpreted to assess the significance and precision of effects.

### Tract Microstructure

For each participant, tract neurite density index (NDI) values were resampled at 100 equidistant nodes (Nodes 0–99) and merged with the relevant biomarker data. At every node, we fitted a linear regression to examine how PNMI predicted node NDI, controlling for ICV and maternal education, and recorded the t-statistic and two tailed, uncorrected p-value. Since running 100 parallel node-wise tests inflates the family-wise error rate, we applied a non-parametric cluster-based permutation correction (49–51). Nodewise linear regressions were first performed at each node, and nodes exceeding an uncorrected threshold of p < .05 were grouped into clusters based on anatomical adjacency. Permutation testing was then used to identify a minimum cluster size required for statistical inference. Specifically, participant labels were randomly permuted 1,000 times, and for each permutation the same nodewise analyses and clustering procedure were repeated. From each permutation, the size of the largest cluster was recorded to form a null distribution of maximum cluster sizes. The 95^th^ percentile of this distribution was taken as the critical cluster-extend threshold, controlling the family wise error rate at α = .05. Only clusters whose size exceeded this threshold were considered significant and assigned a permutation-based cluster-level p value (p_cluster). For every PNMI biomarker, we report the size of each significant cluster, its p_cluster, its direction (positive or negative), the mean t statistic, and regression coefficient within the cluster. This non-parametric, cluster-based procedure makes no distributional assumptions and is therefore appropriate given that the node-wise residuals violated normality.

## Results

### Descriptives and Bivariate Correlations

Demographic information is provided in Table 1, and bivariate correlations for T1 and T2 PNMI are reported in Supplementary Tables 1 and 2, respectively.

### Prenatal Maternal Inflammation and Tract Macrostructure

Next, we conducted hierarchical linear regression analyses to examine whether offspring tract macrostructure between the VTA and limbic regions were associated with PNMI biomarkers (Figure 1). Each biomarker was tested in a separate model, and all models included intracranial volume (ICV) and maternal education level as covariates in Step 1. Bootstrapped parameter estimates and confidence intervals are presented in Table 2. Of T1 PNMI, higher maternal concentrations of soluble TNF receptor II (sTNF-RII) were associated with significantly lower macrostructure in the right VTA–hippocampus (*b* = -0.671, 95% CI [-1.37, - 0.02], ΔR² = 0.049). and right VTA–limbic striatum tracts (*b* = -1.226, 95% CI [-2.71, -0.24], ΔR² = 0.070).

**Figure 1.**
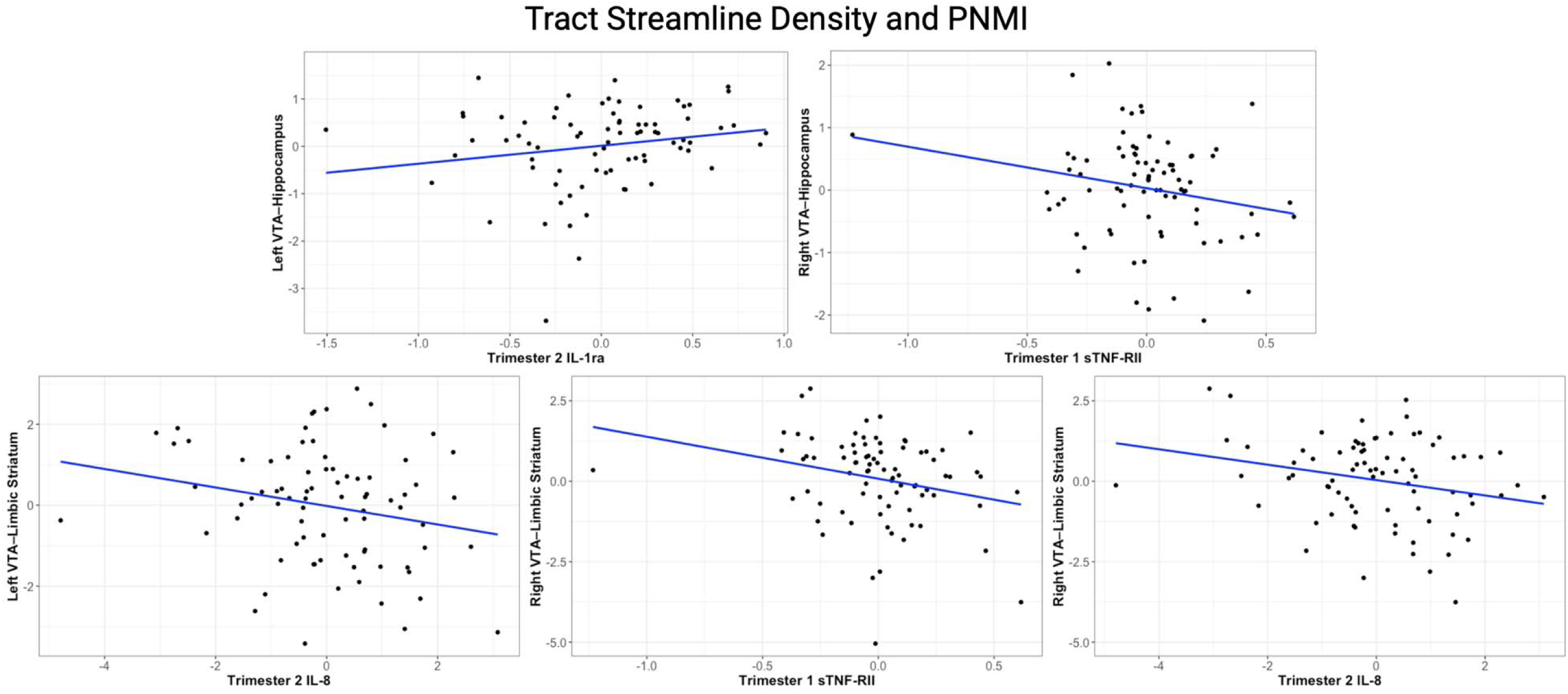
Relationship between maternal serum inflammatory cytokines and offspring tract macrostructure in late midlife. The x-axis represents residualized inflammatory cytokine levels (natural-log-transformed pg/mL) after adjusting for intracranial volume and maternal education. The y-axis represents residualized tract streamline counts (natural-log transformed), controlling for the same covariates.

At T2, higher levels of IL-8 were associated with lower macrostructure in the left (*b* = - 0.248, 95% CI [-0.49, -0.01], ΔR² = 0.055) and right VTA–limbic striatum pathway (*b* = -0.247, 95% CI [-0.47, -0.06], ΔR² = 0.061). Additionally, higher second trimester IL-1ra concentrations were associated with higher macrostructure in the left VTA–hippocampus pathway (*b* = 0.416, 95% CI [0.02, 0.90], ΔR² = 0.046). No other biomarkers were significantly associated with macrostructure in any of the tracts examined.

### Prenatal Maternal Inflammation and Tract Microstructure

All node-wise permutation testing models with at least one significant cluster (2 or more consecutive nodes associated with a cytokine) are reported in Table 3. We identified one significant node cluster that survived cluster correction within each hemisphere of the VTA-limbic striatum and VTA-hippocampus tracts. Higher levels of T1 IL-8 were associated with lower neurite density in the left VTA-hippocampus tract, within the lateral portion of the tract, closer to the hippocampus (nodes 42-84) (*b* = -5.107, SE = 2.224, p = 0.014). Higher levels of T2 IL-6 were associated with higher neurite density in the right VTA-hippocampus tract in a lateral cluster close to the hippocampus (nodes 64-96) (*b* = 3.247, SE = 1.017, p = 0.030), see Figure 2. In addition, higher levels of T1 IL-8 were associated with reduced neurite density within posterior node clusters in the left and right VTA-limbic striatum tracts, closer to the VTA (nodes 12-45, *b* = -4.939, SE = 2.121, p = 0.030; nodes 0-61, *b* = -6.585, SE = 2.339, p = 0.004, respectively), see figure 3.

**Figure 2.**
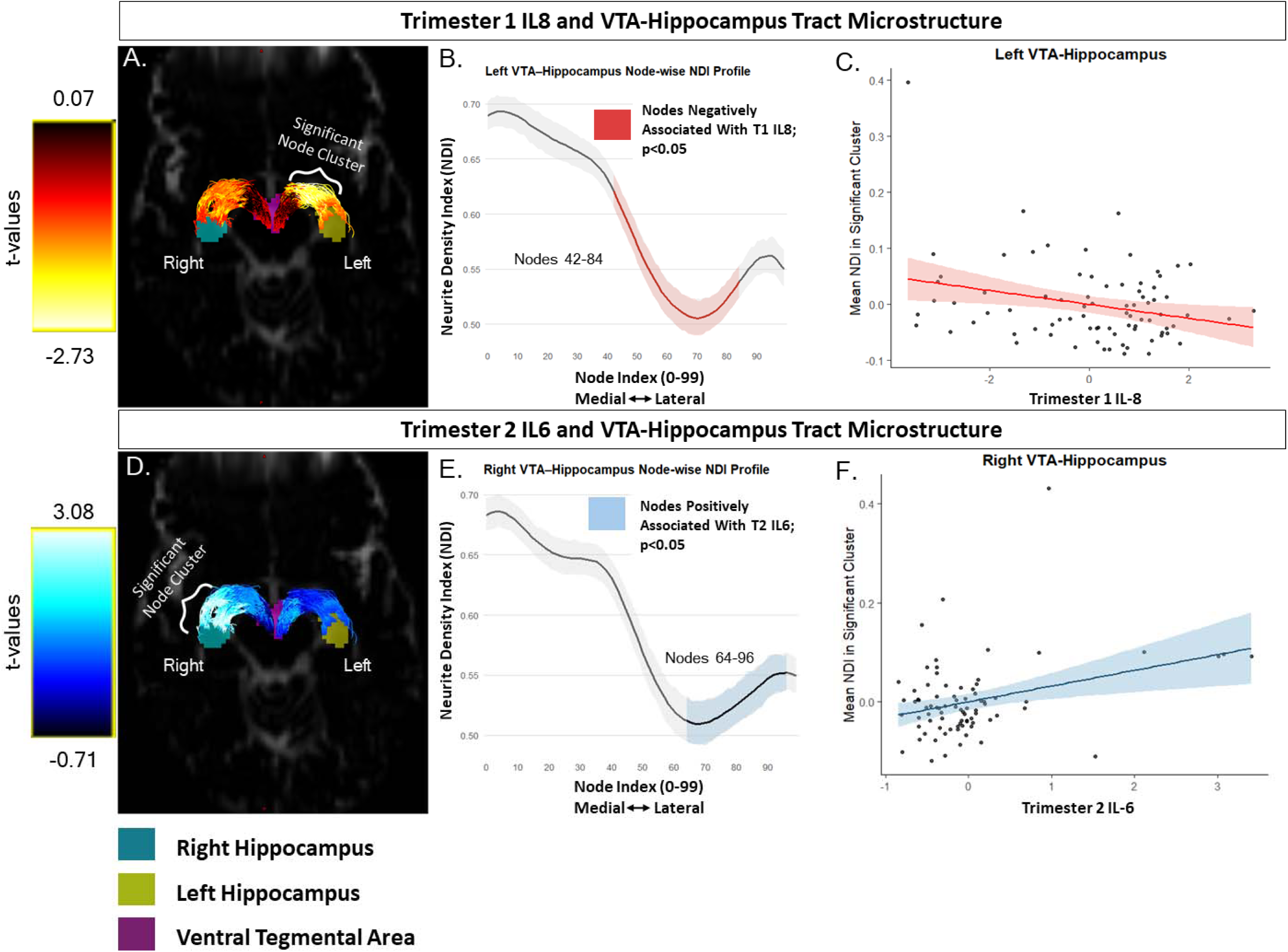
Associations between VTA-Hippocampus tract microstructure (both left and right hemispheres) and prenatal maternal inflammation biomarkers, controlling for intracranial volume and maternal education. ***A,*** significance levels of the relationship between prenatal exposure to IL-8 in the first trimester and neurite density each node along the Left VTA-Hippocampus tract (100 nodes). ***B,*** Node-wise neurite density index (NDI) profile along the Left VTA-Hippocampus tract. The x-axis represents node indicides from medial (0) to lateral (99) along the tract. The y-axis shows the average NDI. The nodes shaded in red (42-84) showed a significant negative association with first trimester IL-8 exposure (p<0.05). Shaded area around the curve represents the 95% confidence interval. ***C,*** The relationship between trimester 1 IL-8 and mean NDI within the significant cluster (nodes 42-84). The x-axis represents residualized trimester 1 IL-8 (natural-log-transformed pg/mL). The y-axis represents residualized mean NDI values. The solid red line indicates the linear regression fit, the light red shaded area represents the 95% confidence interval. ***D,*** significance levels of the relationship between prenatal exposure to IL-6 in the second trimester and neurite density each node along the right VTA-Hippocampus tract (100 nodes). ***E,*** Node-wise neurite density index (NDI) profile along the Right VTA-Hippocampus tract. The x-axis represents node indicides from medial (0) to lateral (99) along the tract. The y-axis shows the average NDI. The nodes shaded in blue (64-96) showed a significant positive association with second trimester IL-6 exposure (p<0.05). Shaded area around the curve represents the 95% confidence interval. ***F,*** The relationship between trimester 2 IL-6 and mean NDI within the significant cluster (nodes 64-96). The x-axis represents residualized trimester 2 IL-6 (natural-log-transformed pg/mL). The y-axis represents residualized mean NDI values. The solid blue line indicates the linear regression fit, the light blue shaded area represents the 95% confidence interval.

**Figure 3.**
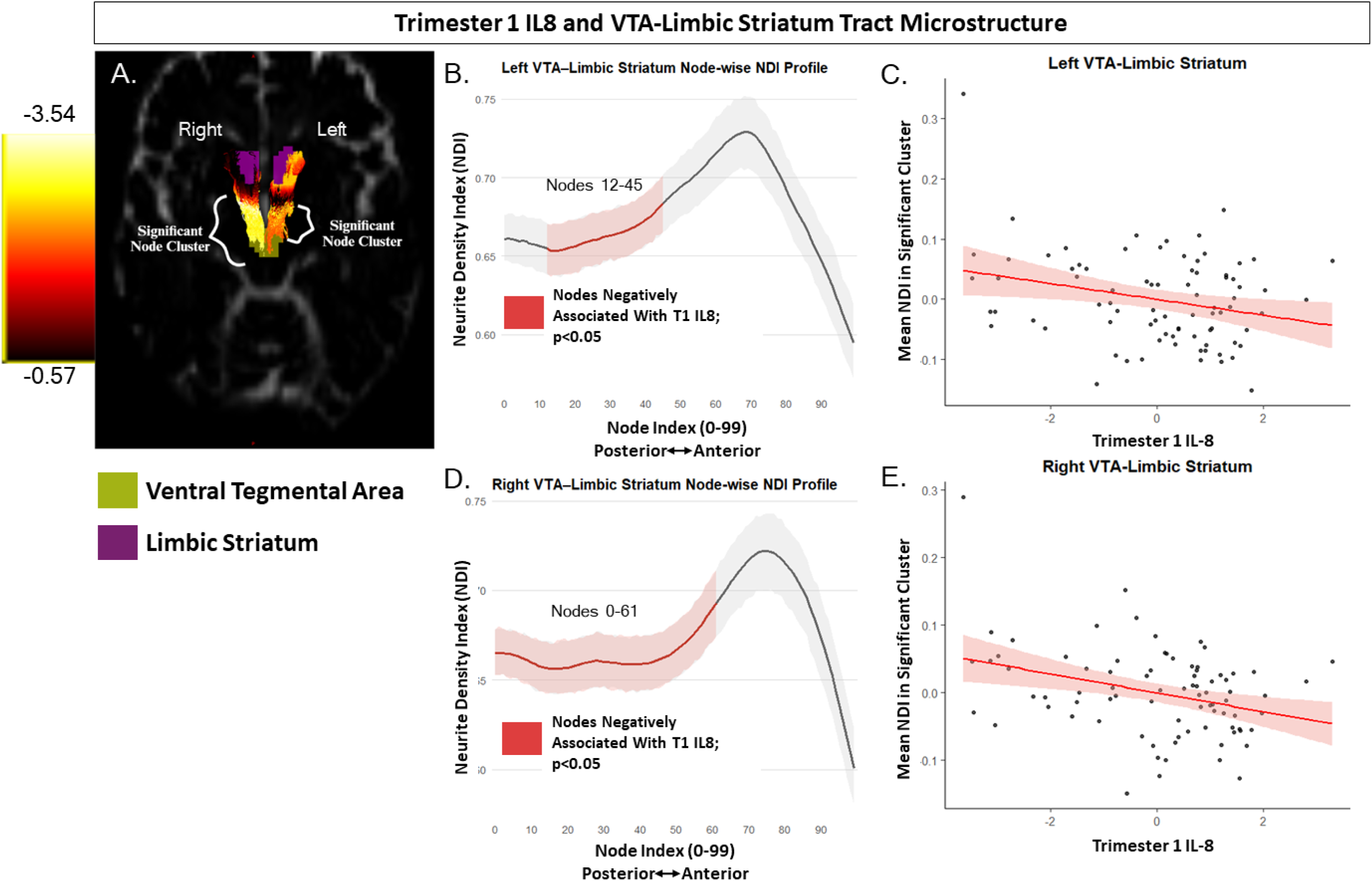
Associations between VTA-Limbic Striatum tract microstructure (both left and right hemispheres) and prenatal maternal inflammation biomarkers, controlling for intracranial volume and maternal education. ***A,*** significance levels of the relationship between prenatal exposure to IL-8 in the first trimester and neurite density each node along both hemispheres of the VTA-Limbic Striatum tract (100 nodes). ***B,*** Node-wise neurite density index (NDI) profile along the Left VTA-Limbic Striatum tract. The x-axis represents node indices from posterior (0) to anterior (99) along the tract. The y-axis shows the average NDI. The nodes shaded in red (12-45) showed a significant negative association with first trimester IL-8 exposure (p<0.05). Shaded area around the curve represents the 95% confidence interval. ***C,*** The relationship between trimester 1 IL-8 and mean NDI within the significant cluster (nodes 12-45). The x-axis represents residualized trimester 1 IL-8 (natural-log-transformed pg/mL). The y-axis represents residualized mean NDI values. The solid red line indicates the linear regression fit, the light red shaded area represents the 95% confidence interval. ***D,*** Node-wise neurite density index (NDI) profile along the Right VTA-Limbic Striatum tract. The x-axis represents node indices from posterior (0) to anterior (99) along the tract. The y-axis shows the average NDI. The nodes shaded in red (0-61) showed a significant negative association with first trimester IL-8 exposure (p<0.05). Shaded area around the curve represents the 95% confidence interval. ***E,*** The relationship between trimester 1 IL-8 and mean NDI within the significant cluster (nodes 0-61). The x-axis represents residualized trimester 1 IL-8 (natural-log-transformed pg/mL). The y-axis represents residualized mean NDI values. The solid red line indicates the linear regression fit, the light red shaded area represents the 95% confidence interval.

**Table 3.**
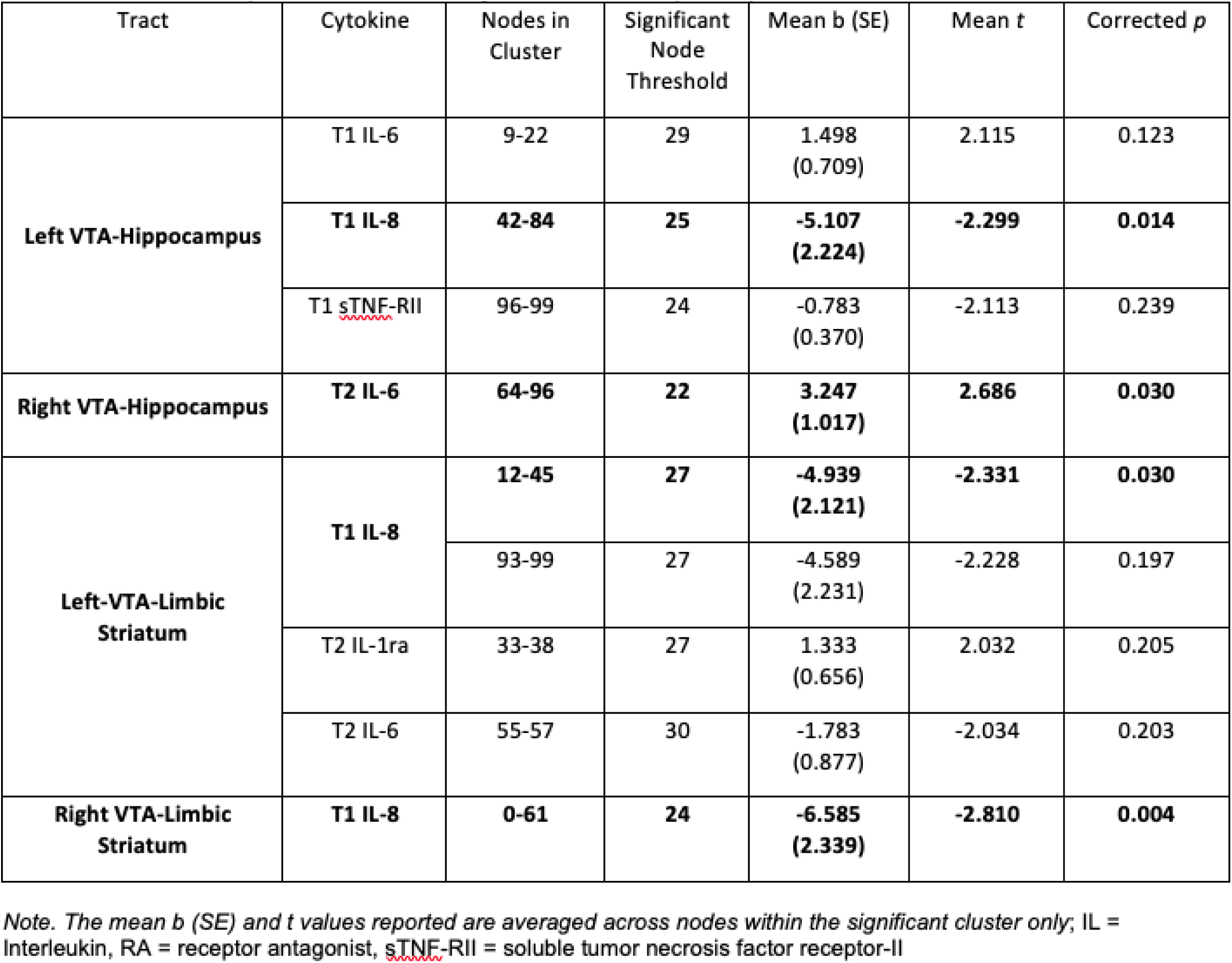
Results of node-wise permutation testing linear regression analyses for prenatal maternal inflammation and offspring tract neurite density index, controlling for intracranial volume and maternal education. Only models that emerged with at least one cluster (2 or more consecutively associated nodes) are reported.

## Discussion

In the current study, we found that elevated PNMI was associated with bidirectional differences in structural connectivity of mesolimbic tracts in offspring, nearly 60 years later. To our knowledge, this is the first study in humans to find an association between PNMI and mesolimbic circuit structure later in life. We measured two distinct indices of tract integrity: macrostructure, derived from probabilistic tractography, and microstructure (NDI), derived from the NODDI model and linked PNMI to changes in both measures of mesolimbic structural connectivity. Our findings partially supported our directional hypotheses, with elevated PNMI being associated with reduced micro- and macrostructure in mesolimbic pathways. However, a couple of cytokines exhibited associations in the opposite direction, suggesting that prenatal inflammatory exposures may exert heterogenous and regionally specific effects on mesolimbic circuitry. Together, these findings suggest that prenatal inflammation is associated with long-lasting changes in offspring mesolimbic circuits. The sections below integrate these findings with prior literature, organized by cytokine to clarify potential mechanistic pathways.

### IL-8

Higher prenatal IL-8 was associated with reduced offspring macro- and microstructure, but with differences in timing. Specifically, higher prenatal IL-8 during the second trimester was associated with reduced macrostructure in the right VTA-hippocampus and VTA-limbic striatum tracts of offspring, while higher IL-8 during the first trimester was associated with reduced neurite density along the left VTA–hippocampus tract (closer to the hippocampus) and in both the left and right VTA–limbic striatum tracts of offspring (closer to the VTA). These findings suggest that IL-8 may have widespread and temporally specific effects on the development of mesolimbic circuitry. Notably, this aligns with findings from our previous study with this cohort, where first trimester IL-8 was linked to reduced hippocampal neurite density in offspring during late middle age (16), further supporting the hippocampus as a site of early vulnerability to inflammatory signals and implicating mesolimbic neuromodulatory systems. Moreover, these results comport with findings by (10), who found elevated prenatal IL-8 was associated with reductions in offspring entorhinal cortex volume and increased cerebrospinal fluid among cases with schizophrenia spectrum disorders. The entorhinal cortex plays a key role in gating information flow into the hippocampus and is deeply integrated into mesolimbic circuitry (52).

These IL-8-related disruptions may therefore reflect early impairments in a broader cortico-limbic network that regulates memory, emotion, and novelty processing. Importantly, such circuit-level alterations are consistent with the theoretical framework put forth by (18), which posits a hippocampal-VTA circuit that coordinates novelty detection and dopaminergic signaling to guide motivated behavior. In this model, hippocampal input to the limbic striatum and VTA drives a feedback system critical for salience detection, memory encoding, and behavioral flexibility (17,53). Disruption to this circuit by prenatal inflammation may have lasting consequences for the regulation of affect, motivation, and learning.

### IL-6

Contrary to our hypotheses, we found that higher IL-6 during the second trimester was associated with *increased* neurite density along the VTA–hippocampus tract but without accompanying changes in macrostructure. The direction of the effects is puzzling but not entirely inconsistent with the prior literature. For instance, consistent with the present study, a rodent study found IL-6 injections in pregnant mice at gestation day 15, analogous to late T2 or early T3 in humans, was linked to hyperconnectivity within the offspring brain in adulthood, particularly in hippocampal regions (54). One possibility for our findings is that increased neurite density along the VTA-hippocampus pathway may reflect altered developmental refinement of mesolimbic circuitry. Neurodevelopment involves early overproduction of neurites followed by activity-dependent pruning, and IL-6 plays a central role in immune signaling and microglial activity that regulates synaptic loss and remodeling (55). Elevated T2 IL-6 may therefore disrupt normative axonal refinement within VTA-hippocampus projections, leading to aberrant pruning and maladaptive circuit organization rather than enhanced structural integrity. The unilateral nature of this association should be interpreted cautiously. Subcortical regions, including the hippocampus, have been shown to have structural asymmetry across hemispheres (56), and the localization of IL-6 associated nodes closer to the hippocampus may reflect such asymmetry.

However, the present study was not designed to test hemispheric differences directly, and the observed asymmetry may reflect statistical variability rather than true biological lateralization.

Additionally, a previous study from our lab with the HBP cohort found that increased prenatal IL-6 during the first trimester was associated with reduced hippocampal neurite density (16), a finding in the opposite direction of the present findings. An explanation for the inconsistency between our prior trimester 1 and present trimester 2 findings is that the differences in directionality and timing may reflect a developmental shift in vulnerability, where early gestational IL-6 exposure disrupts hippocampal gray matter microstructure directly, while later exposure may alter connectivity patterns (axons) between the hippocampus and neuromodulatory regions like the VTA. These findings underscore the importance of considering timing-specific effects of PNMI, as the hippocampus undergoes distinct maturational processes across trimesters that may render it differentially sensitive to inflammatory insults (57).

### IL-1ra

Interestingly and also contrary to our hypotheses, we found that prenatal IL-1ra during the second trimester was positively associated with increased tract density from VTA to hippocampus in the left hemisphere of offspring. The mechanisms through which elevations in a pro-inflammatory marker is associated with elevated tract macrostructure remains unclear.

However, from a developmental perspective, it could be that elevated T2 IL-1ra may influence the large-scale organization of this long-range mesolimbic projection. Unlike the T2 IL-6 findings, which were localized to neurite density within the tract, the association with IL-1ra was observed at the level of overall tract density, suggesting potential alterations in pathway formation or tract-level organization. The second trimester and onwards is characterized by active axonal extension, pathfinding, establishment of neurotransmitter systems, and assembly of long-range circuits (58). Inflammatory perturbation during this window may therefore affect the establishment or stabilization of long-range projections between the VTA and hippocampus, potentially increasing tract density through altered axonal growth or reduced elimination of developing projections. As with the IL-6 findings, these results may reflect altered developmental refinement rather than enhanced structural capacity. From an adaptive perspective exposure to prenatal maternal immune activation during this period may serve as a signal that calibrates the developing mesolimbic system to anticipated postnatal environmental conditions, particularly those involving uncertainty or stress. In this context, the observed changes in structural connectivity may reflect differences in the refinement of developing circuits.

Additionally, the hemispheric specificity of this association should be interpreted cautiously, and replication will be necessary before attributing the findings to stable biological asymmetry.

### sTNF-RII

Consistent with our hypotheses, we observed a negative relationship between second trimester sTNF-RII and macrostructure of the white matter tract connecting the VTA to the hippocampus in the right hemisphere. Although a previous study found no relationship with sTNF-RII and hippocampal microstructure in this cohort (16), the two studies examine fundamentally different biological targets. Specifically, Mohyee et al., (2025) examined gray matter microstructure within the hippocampus, using NODDI-derived neurite density index to model local dendritic architecture. Their null finding therefore pertains to cellular-scale organization within hippocampal gray matter. In contrast, the present findings relate T2 sTNF-RII to VTA-hippocampal white matter macrostructure, a measure indexing long-range structural connectivity and large-scale communication capacity, suggesting that PNMI may differentially relate to microstructural gray matter versus circuit-level white matter organization. Our findings align with recent studies that found that higher gestational exposure to maternal TNF-α was associated with poorer memory performance and with differences in hippocampal fMRI activation during the performance of memory tasks (15,59). This pattern of findings suggests that the relationship between sTNF-RII and hippocampal dysfunction may be due, in part, to structural alterations within neuromodulatory circuits that influence hippocampal function (18,53,60). These results lay the groundwork for future studies to investigate the role of sTNF-RII on modulatory circuits upstream of the hippocampus that influence its physiology and function.

### Macrostructure vs Microstructure

Our observed differences between macro- and microstructural findings suggest that mesolimbic circuits may be differentially sensitive to exposure to PNMI depending on the biological metric assessed. Tract macrostructure reflects the gross organization of a pathway, capturing features such as the overall volume and geometry of each fiber bundle, whereas microstructural indices provide voxel-wise estimates of tissue composition that are distinct from a tract’s shape and volume (40). Macrostructure was measured using streamline counts from tracts generated using anatomically informed tractography. Given our interest in these two specific tracts, this approach enabled anatomically constrained evaluation of the strength of pathways in their entirety, as compared to prior, more commonly used whole-brain fractional anisotropy measures used in DTI studies (61–63). Studies using macrostructure techniques have been validated against histological fiber tracing methods (64,65) and tractography-derived macrostructure of reward circuits has been associated with various aspects of human behavior such as reward-motivated memory (37), novelty seeking and reward dependance (66), and impulsivity and decision making (37,67–69). Microstructure was modeled using the neurite orientation dispersion and density imaging (NODDI) model. The NODDI model estimates the complexity of neurites (dendrites and axons) using a multi-compartment diffusion model (43). The NODDI model allows for more granular insight into microstructural changes compared to traditional diffusion tensor imaging (DTI) techniques (43) and has been shown to be useful in early detection of neurodegenerative changes (70,71) thereby rendering it suitable for the current study. These complementary approaches allowed us to investigate dissociable aspects of white matter cytoarchitecture and how PNMI may differentially influence macro- and microstructural properties of offspring mesolimbic circuitry The differential results reported here underscore the utility of employing both approaches.

### Mesolimbic Pathways and Mental Health

The VTA is composed primarily of dopaminergic neurons but also some GABAergic and glutamatergic neurons (72,73). Both dopaminergic and GABAergic signaling within the VTA and its projections are developmentally sensitive to early immune perturbations. Preclinical models of maternal immune activation have found that prenatal inflammation exposure alters dopamine neuron development, reduces dopaminergic fiber density in the nucleus accumbens, and impairs mesolimbic dopamine transmission in offspring (31,32,74). Additionally, PNMI has been shown to disrupt GABAergic interneuron maturation and long-range GABAergic projections, particularly within circuits involved in motivation and stress regulation (75–77).

Such alterations are associated with symptoms in disorders such as schizophrenia, depression, and ADHD, including anhedonia, cognitive inflexibility, and affective blunting (78–81). Despite the robust animal literature implicating PNMI in altered dopamine and GABA system development, human studies have only recently begun to map these relationships in vivo. The present findings, which show PNMI-related changes in tract and neurite density along mesolimbic pathways, offer novel evidence that prenatal immune exposure may influence the integrity of these potentially dopaminergic and GABAergic projections in humans well into late middle age. This circuit-level disruption may represent one mechanism linking PNMI to enduring risk for psychiatric symptoms across the lifespan.

Additionally, research has shown that PNMI exerts lasting effects on offspring memory circuitry and cognitive performance (59,82). Moreover, PNMI has been associated with increased risk for schizophrenia and structural alterations within limbic regions (10). Elevated PNMI has also been linked to increased risk for adolescent alcohol use, and substance use disorder diagnoses in late midlife (83,84). Given that mesolimbic circuitry is critically involved in memory, affective regulation, and reward processing, future research should examine whether the present findings extend to psychiatric outcomes such as memory impairment, positive and negative symptoms, and substance use.

### Limitations and Future Directions

The present study has several limitations. First, although this study capitalized on a unique cohort, our sample size only allowed for modest power to detect significant associations. Second, it was beyond the scope of the present study to examine the various sources of maternal inflammation, such as infection, stress, obesity, health conditions (85) all of which may have independent or combined effects on offspring neurodevelopment, thereby limiting our ability to draw conclusions about potential environmental factors that lead to elevated levels of PNMI. Second, although our selection of PNMI biomarkers was guided by *a priori* hypotheses, these markers may only partially reflect the complex immune dynamics of pregnancy, offering limited insight into broader immunological processes that could influence offspring neurodevelopmental trajectories. Third, it is possible that samples may have degraded. Despite the age of the biological samples, rigorous protocols were implemented to preserve their integrity, analyses focused on cytokines typically present at higher concentrations in pregnant serum, and careful reliability procedures were in place to ensure that protein levels were similar to contemporary samples, see (7) for details. Moreover, it is unlikely that sample degradation occurred in a systematic manner that would influence our results. Fourth, our neuroimaging methods do not distinguish cell types. Future studies could employ PET imaging to measure neuromodulator and neurotransmitter concentrations in-vivo (86–88) to determine how prenatal immune perturbations shape signaling of specific neurotransmitters and neuromodulators in mesolimbic circuits.

Last, there is a substantial temporal gap between prenatal inflammatory exposure and assessment of brain structure in late midlife, which constrains inferences about underlying mechanisms. While PNMI may exert long-lasting effects by altering early neurodevelopmental processes such as cell differentiation, axonal guidance, and circuit assembly, our data cannot determine whether the observed brain differences reflect direct prenatal programming or the cumulative influence of postnatal immune processes across the lifespan. However, given the temporal precedence of prenatal immune activation, any downstream immune phenotype in the offspring would most plausibly serve as a moderator or mediator of the observed associations, rather than a confounder or an alternative explanation. Nevertheless, prior preclinical work demonstrates that prenatal immune activation can produce enduring hippocampal alterations even in the absence of persistent offspring inflammation (89), suggesting that prenatal mechanisms alone may be sufficient to generate the long-term structural effects observed in the present study. Nevertheless, future studies incorporating longitudinal immune measures across development will be necessary to disentangle prenatal programming effects from postnatal immune modulation, especially given that prenatal inflammation can induce inflammatory states in offspring (90).

The magnitude of the associations observed between prenatal maternal inflammation and mesolimbic tract macrostructure was small to moderate, with inflammatory biomarkers accounting for approximately 4 to 7 percent of incremental variance beyond covariates. These effect sizes are consistent with prior studies studies linking prenatal inflammatory exposures to later brain structure and function (10,15,16). The small-to-moderate effect sizes observed in the present study may also reflect the influence of numerous environmental, biological, and psychosocial factors that unfold across the lifespan following the prenatal period. Given the late midlife age of the offspring sample, decades of postnatal exposures, including stress, illness, lifestyle factors, and aging-related neurobiological changes, are likely to contribute additional variance in brain structure, thereby attenuating the proportion of variance attributable to prenatal immune exposure alone. Such attenuation is expected in prospective developmental studies with long temporal gaps between exposure and outcome and does not diminish the biological relevance of the observed associations. Notably, similar effect sizes have been reported in developmental cohorts assessed closer to the prenatal period (11), although differences in imaging techniques and white matter tracts examined make direct comparison difficult.

## Conclusion

This study tested participants from a rare longitudinal birth cohort with prospectively collected prenatal blood serum and neuroimaging data acquired among offspring during late middle age. The availability of trimester-specific, serologically defined inflammatory biomarkers enables fine-grained investigation of how distinct periods of prenatal immune activation influence the long-term structure of mesolimbic dopamine pathways. Our study provides a significant development in our understanding of how PNMI has lasting impacts on mesolimbic circuit structure in offspring at late midlife. Our findings reveal an association between elevated maternal inflammation during pregnancy and structural alterations in key mesolimbic tracts, suggesting that prenatal immune programming may shape neural circuitry linked to reward processing, memory, and affective regulation. These results provide insight into how prenatal programming might contribute to aging-related changes in the brain, emphasizing the need for further exploration of the underlying biological mechanisms that may contribute to psychiatric and age-related neurobehavioral changes.

## Notes

### Competing Interest Statement

The authors have declared no competing interest.

### Funding

- National Institute of Mental Health: R01 MH096478, MH118545
- Eunice Kennedy Shriver National Institute of Child Health and Human Development: HHSN275201100020C

