## supplementary information for "Prenatal Maternal Inflammation Is Associated with Altered Offspring Mesolimbic White Matter Circuitry Observed in Late Midlife"

**Prenatal Maternal Inflammation (PNMI) Data**

Data were available for PNMI biomarker concentrations determined in archived maternal sera from a subset of pregnant mothers in the parent CHDS study (n=737), during T1 (mean gestational age in weeks 12.1 ± 2.6) and/or T2 (mean gestational age in weeks 23.7 ± 3.0), as previously described (1–3). The biomarkers were interleukin-6 (IL-6), IL-8, IL-1 receptor antagonist (IL-1ra), and soluble tumor necrosis factor receptor-II (sTNF-RII). Assays utilized high sensitivity (IL-6, IL-8) and regular sensitivity (IL-1ra, sTNF-RII) ELISAs (R&D Systems), with detectable concentrations on all samples for all biomarkers, and inter- and intra-assay correlation coefficients of <9% and <4%, respectively. Both IL-1ra and sTNF-RII were chosen as proxies for IL-1β and TNF-⍺ activity, respectively, as they are found at higher concentrations and are more consistently detectable in serum. For more details on the inflammatory biomarker assays, refer to supplement in (1).

### **MRI Data acquisition**

The MRI acquisition parameters are also described in (4). For consistency and clarity, we replicate the description here. MRI scans were acquired on a 3 Tesla Siemens MAGNETOM Trio scanner using a 32-channel head coil at the University of California, Berkeley, Henry H. Wheeler Jr. Brain Imaging Center. A T1-weighted structural image was acquired using multi-echo, magnetization-prepared, 180-degree radio-frequency pulses and rapid gradient-echo (MEMPRAGE) sampling (TR=2,530.0 ms, TE=1.64-7.22 ms, FOV=256.0 mm, flip angle=7°, slice thickness=1.0 mm, slices per slab=176.0, voxel size=1.0 × 1.0 × 1.0 mm, matrix size=176 × 256 × 256 voxels, scan time [min]=6:03). Diffusion-weighted images were collected along 111 directions using a multi-shell sequence (b = 1000 s/mm2 and b = 2000 s/mm2) and sensitivity encoding image reconstruction (SENSE) (TR=3,400.0 ms, TE=94.8 ms, FOV=210 mm, flip angle=78°, slice thickness=2.0 mm, slices=69, voxel size=2.0 × 2.0 × 2.0 mm, matrix size=104 × 104 × 69 voxels), scan time [min]=6:31). Additional b0 (no diffusion weighting) images were collected in the reverse phase encoding direction to allow for the estimation of susceptibility-induced distortions in the preprocessing of the diffusion scans.

**Diffusion-weighted imaging (DWI) Preprocessing**

Diffusion-weighted scans were denoised using dwidenoise from MRtrix3 Version 3.0.2 (5,6). Susceptibility-induced distortions were then estimated according to (7) using the command topup from FSL Version 6.0.5.1 (8). Eddy current distortion correction and susceptibility-induced distortion correction was run using FSL’s eddy with the slice-to-volume motion correction and outlier replacement flags (9–11). The corrected images were then visually assessed for remaining image quality issues before NODDI metrics were estimated.

#### **Probabilistic Tractography**

Probabilistic tractography was performed in MRTrix3 (<https://www.mrtrix.org/>). Voxel-wise response functions were estimated for each principal tissue compartment, (e.g., gray matter, white matter, and CSF) using the *dhollander* algorithm (12,13). Multitissue orientation distribution functions for each macroscopic tissue type were fitted using the *msmt_csd* argument in the *dwi2fod* command. This approach capitalized on the distinct diffusion properties of different tissue types measured by the multi-shell, high angular resolution diffusion imaging (HARDI) sequence used in the current study.

Probabilistic tractography was performed using the iFOD2 algorithm using the *tckgen* command in subjects’ native space. All tractographies seeded in the VTA and were performed unidirectionally, with the upstream target (hippocampus or limbic striatum) designated as both the inclusion and stopping criteria. To allow inferences about tract density (macrostructure), we seeded a fixed number of random streamline attempts per voxel in the seed mask. To control for individual variation in seed size, the lowest volumetric occurrence of the VTA within the cohort was determined. We found the smallest size of the VTA in any subject to have a volume of 276 voxels. Using 5,000 seeding attempts per voxel as the default selection criteria, we calculated the total number of viable seeds for all tractographies by multiplying 276 voxels by 5,000 seed attempts per voxels, resulting in 1,380,000 total seeds for each tractography; an approach that was similar to prior tractography work (14). For each tract, we excluded either the left or right hemisphere to ensure we only captured streamlines that projected to the target region ipsilaterally for all subjects.

**Supplementary Table 1.**

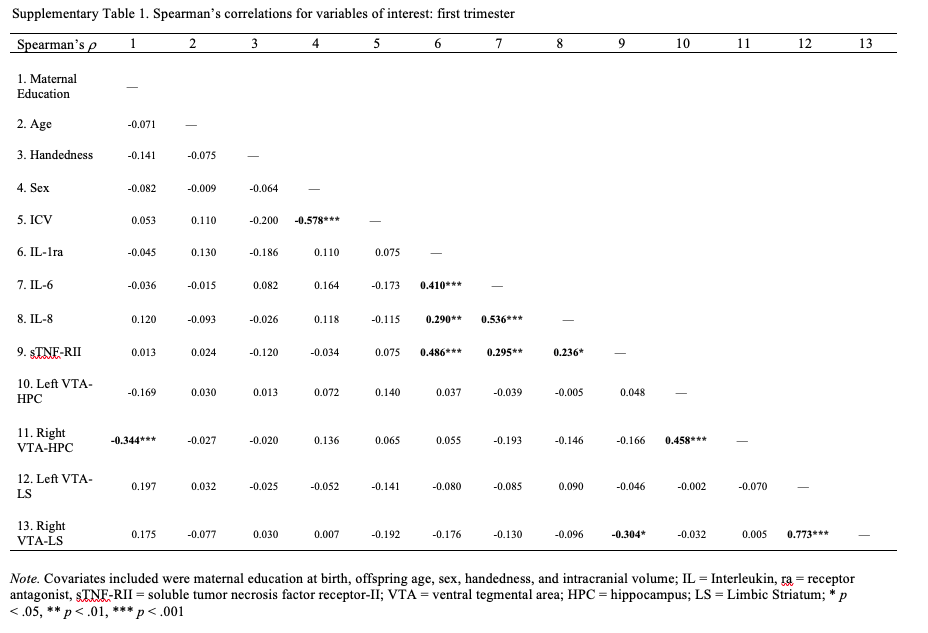

**Supplementary Table 2.**

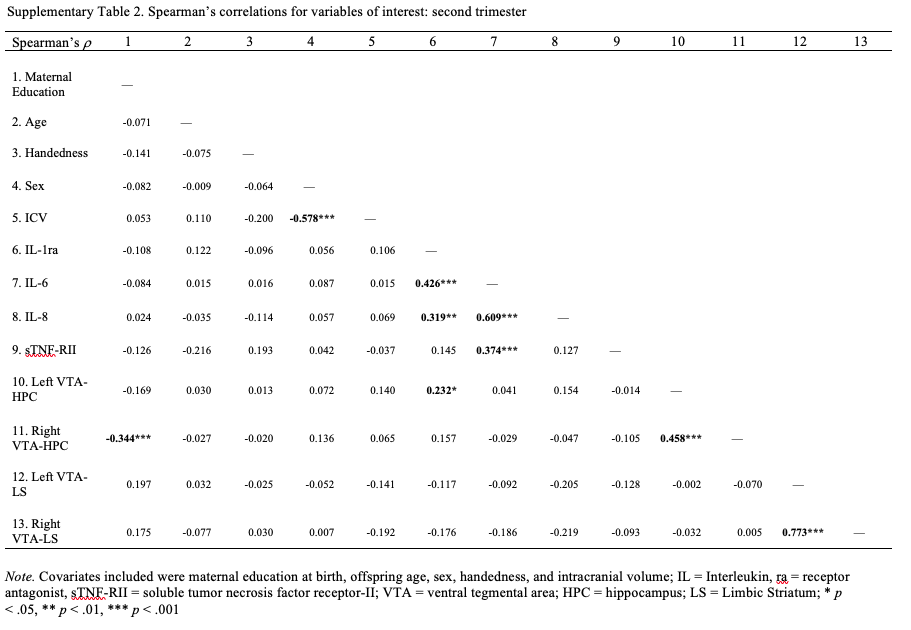

**Supplementary Figure 1.**

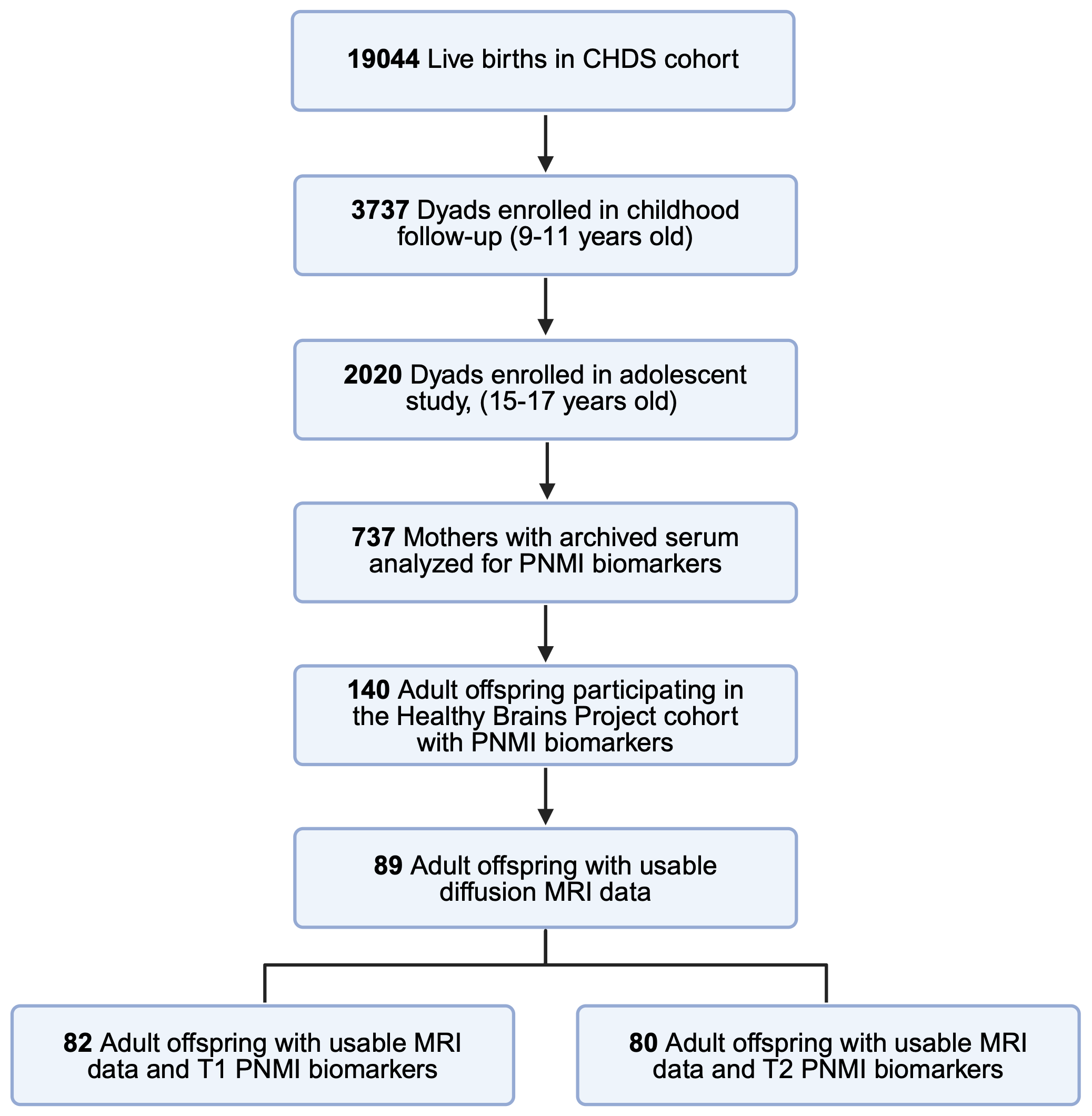

Participants retained in subsamples of the Child Health and Development Studies (CHDS) and Healthy Brain Project (HBP), to yield the final analytic sample for the present study. T1 = first trimester of pregnancy; T2 = second trimester of pregnancy.
